# Mechanism of molnupiravir-induced SARS-CoV-2 mutagenesis

**DOI:** 10.1101/2021.05.11.443555

**Authors:** Florian Kabinger, Carina Stiller, Jana Schmitzová, Christian Dienemann, Hauke S. Hillen, Claudia Höbartner, Patrick Cramer

## Abstract

Molnupiravir is an orally available antiviral drug candidate that is in phase III trials for the treatment of COVID-19 patients^1,2^. Molnupiravir increases the frequency of viral RNA mutations^3,4^ and impairs SARS-CoV-2 replication in animal models^4-6^ and in patients^2^. Here we establish the molecular mechanisms that underlie molnupiravir-induced RNA mutagenesis by the RNA-dependent RNA polymerase (RdRp) of the coronavirus SARS-CoV-2. Biochemical assays show that the RdRp readily uses the active form of molnupiravir, β-D-N4-hydroxycytidine (NHC) triphosphate, as a substrate instead of CTP or UTP. Incorporation of NHC monophosphate into nascent RNA does not impair further RdRp progression. When the RdRp uses the resulting RNA as a template, NHC directs incorporation of either G or A, leading to mutated RNA products. Structural analysis of RdRp-RNA complexes containing mutagenesis products shows that NHC can form stable base pairs with either G or A in the RdRp active center, explaining how the polymerase escapes proofreading and synthesizes mutated RNA. This two-step mutagenesis mechanism likely applies to various viral polymerases and can explain the broad-spectrum antiviral activity of molnupiravir.

---

Coronaviruses use an RNA-dependent RNA polymerase (RdRp) for the replication and transcription of their RNA genome^7-11^. The RdRp is an important target for the development of antiviral drugs against coronaviruses^12-15^. Structures of the RdRp were reported for SARS-CoV-1^16^ and for SARS-CoV-2^17-21^ and provided insights into the mechanisms of RNA-dependent RNA synthesis^22^. The structures also enable mechanistic studies that can rationalize the molecular processes underlying the antiviral activity of compounds targeting the RdRp.

Antiviral drugs often target viral polymerases and function as nucleoside analogs that terminate RNA chain elongation. However, such chain-terminating antivirals are generally not effective against SARS-CoV-2 because coronaviruses carry an exonucleolytic proofreading activity that can remove misincorporated nucleotides from the nascent RNA 3’-end^23-25^. The nucleoside analogue remdesivir can circumvent proofreading because its incorporation does not terminate elongation but only stalls RdRp after the addition of three more nucleotides^19,26-29^. Remdesivir was the first FDA-approved drug for the treatment of COVID-19 patients^30-33^, but its effectiveness is disputed^34^, emphasizing the need to develop new antiviral drugs.

Another promising drug candidate for the treatment of COVID-19 patients is molnupiravir (or EIDD-2801), which also targets the RdRp of SARS-CoV-2. Molnupiravir is an isopropylester prodrug of the nucleoside analogue β-D-N4-hydroxycytidine (NHC or EIDD-1931)^3,35^. Molnupiravir interferes with the replication of various viruses^3,35-40^ including SARS-CoV-2^4,41^. Molnupiravir inhibits SARS-CoV-2 replication in human lung tissue^5^, blocks SARS-CoV-2 transmission in ferrets^6^, and reduces SARS-CoV-2 RNA in patients^2^. In contrast to approved drugs such as remdesivir that are administered by infusion, molnupiravir is orally available. Molnupiravir has been tested in phase I trials^42^ for safety, tolerability and pharmacokinetics, and phase II/III studies are currently ongoing^2^ (NCT04405739, NCT04405570, NCT04575597). Available data suggest that molnupiravir acts as a mutagenizing agent that causes an ‘error catastrophe’ during viral replication^3,4,43^. Indeed, NHC can introduce mutations into viral RNA, as shown for Venezuelan equine encephalitis virus^44^. Also, sequencing of influenza virus populations indicated that NHC caused G-to-A and C-to-U transitions in viral RNA^3^, and the same transitions were found for SARS-CoV-2^4^.

Despite this progress, a systematic biochemical and structural analysis of molnupiravir- or NHC-induced RNA mutagenesis by viral RNA polymerases is lacking. Here we quantify the effects of molnupiravir/NHC on the RNA synthesis activity of SARS-CoV-2 RdRp using a purified biochemical system and defined synthetic RNAs. Together with structural analysis we establish the molecular mechanism of molnupiravir-induced RNA mutagenesis. Our results provide detailed insights into the mechanism of action of molnupiravir, which is entirely distinct from that of remdesivir or chain-terminating nucleoside analogues.

## RESULTS

### SARS-CoV-2 RdRp readily incorporates NHC into RNA

We first tested whether purified SARS-CoV-2 RdRp can use the active form of molnupiravir, NHC triphosphate (‘MTP’, **Fig. 1a, b**), as a substrate for RNA synthesis. We conducted RNA elongation assays in a defined biochemical system using recombinant RdRp and synthetic RNA template-product duplexes (Methods). We used four different RNA duplexes that differed at position +1 of the template strand (**Supplementary Table 1**), which directs binding of the incoming nucleoside triphosphate (NTP) substrate (**Fig. 1c**). The RNA product strand contained a fluorescent label at its 5’-end that allowed us to monitor and quantify RNA elongation.

**Figure 1.**
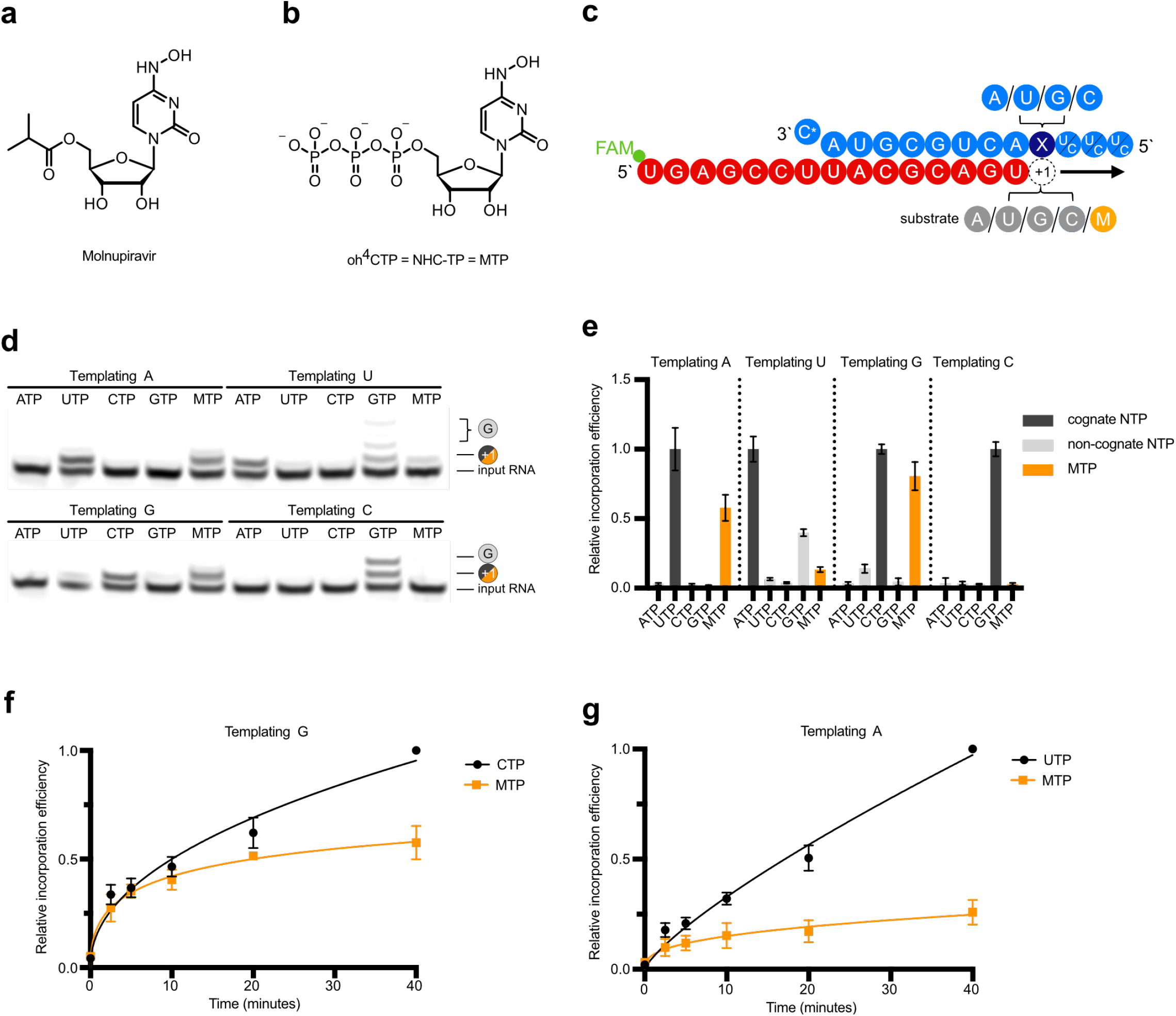
RdRp incorporates NHC opposite of G and A in the template. **a**, Chemical structure of molnupiravir. **b**, Chemical structure of NHC triphosphate (MTP). **c**, RNA template-product duplex. The direction of RNA extension is shown. Color of depicted circles indicates experimental design: blue, RNA template strand; dark blue, +1 templating nucleotide; red, RNA product strand; gray, NTP substrate; orange: MTP. The 5’-end of the RNA product contains a FAM fluorescent label. C* in 5’ end of the template denotes dideoxy-C (ddC). **d**, NHC monophosphate is incorporated into growing RNA instead of C or U when G or A are present in the template +1 position. **e**, Quantification of nucleotide incorporation efficiency relative to the cognate NTP (dark grey). Non-cognate NTPs and MTP are depicted in light grey and orange, respectively. Standard deviations are shown. Source data are provided as a Source Data file. **f**, Quantification of time-dependent M incorporation opposite a templating G residue after triplicate measurements. Incorporation efficiency is calculated relative to cognate C incorporation. Standard deviations are shown. **g**, Quantification of time-dependent M incorporation opposite a templating A residue after triplicate measurements. Incorporation efficiency is calculated relative to cognate U incorporation. Standard deviations are shown.

When the nucleotides G or A were present at the RNA template position +1, NHC monophosphate (‘M’) was readily incorporated instead of C or U, respectively (**Fig. 1d, e**). Time-dependent RNA elongation experiments showed that M was slightly less efficiently incorporated as the cognate nucleotide C, but less efficiently than the cognate nucleotide U (**Fig. 1f, g**). These results can be explained by base pairing of an incoming MTP substrate with either G or A in the RNA template strand. Consistent with this model, NHC adopts different tautomeric forms^45^ that were predicted to allow for base pairing with either G or A^46^.

### RdRp does not stall after NHC incorporation

We next tested whether the incorporation of NHC monophosphate (M) into nascent RNA interferes with further RNA extension. We first conducted RNA elongation assays with a scaffold that allowed for RNA extension by four nucleotides (nt) (**Fig. 2a**). We observed that incorporation of M instead of the cognate U or C did not prevent incorporation of three subsequent nucleotides (**Fig. 2b**). Further, we tested RNA extension with a scaffold that allowed for the incorporation of 11 nt (**Fig. 2c**). Also in this case the RdRp reached the end of the template when UTP or CTP were replaced by MTP, although again incorporation of M instead of U was less efficient that incorporation of M instead of C (**Fig. 2d**). These results demonstrate that M incorporation into nascent RNA does not prevent further RNA elongation. Thus, longer RNA products containing M nucleotides may be synthesized by the RdRp in the presence of MTP. This posed the question what happens when M-containing RNA is used as a template in a second step.

**Figure 2.**
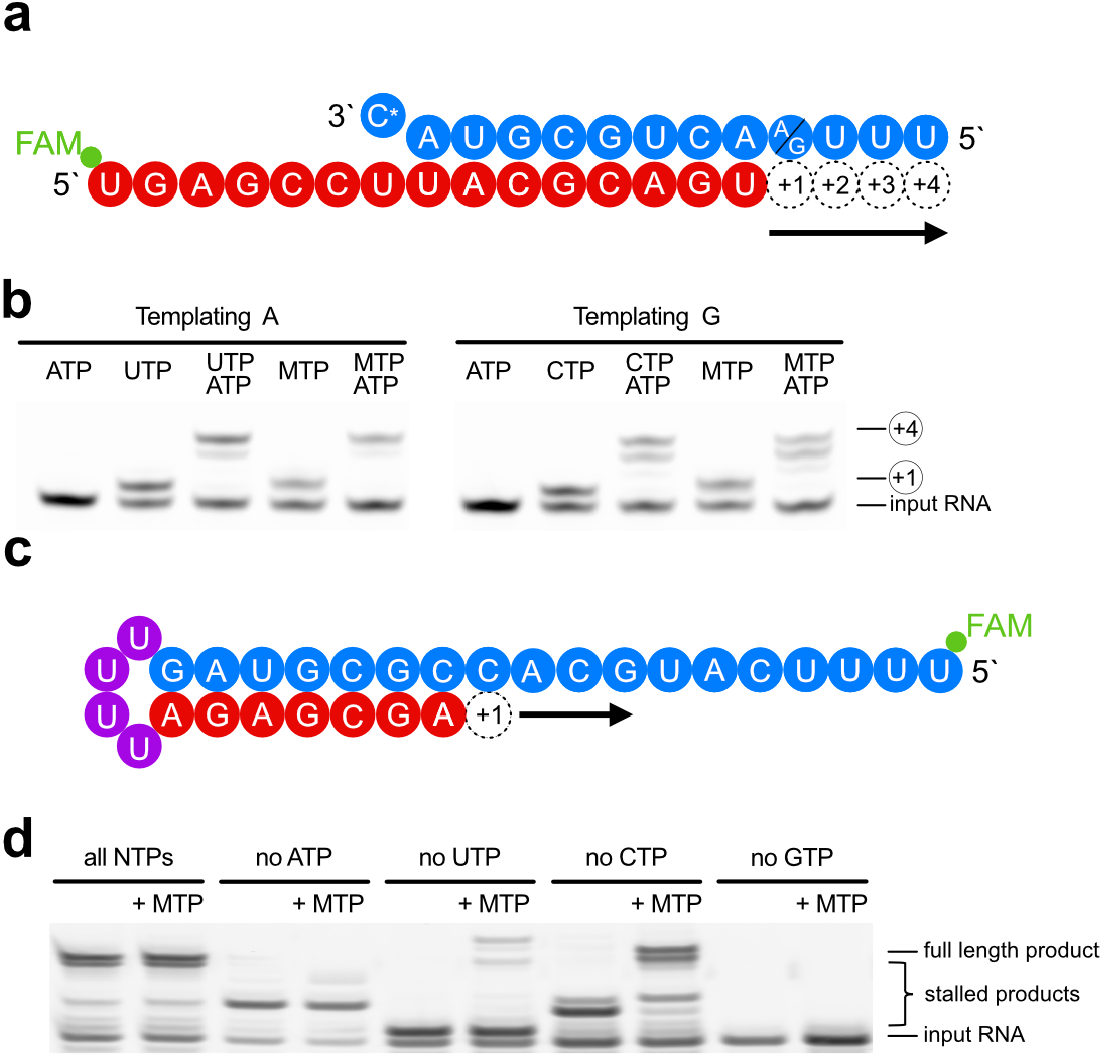
NHC incorporation does not stall SARS-CoV-2 RdRp. **a**, RNA template-product duplex (as in Fig. 1c) allows for RNA extension by four nucleotides. The direction of RNA extension is indicated. The 5’-end of the RNA product contains a FAM fluorescent label. C* indicates dideoxy-C (ddC). **b**, RNA elongation to the end of the template in panel a is possible when MTP replaces either CTP or UTP in the presence of ATP. Source data are provided as a Source Data file. **c**, RNA template-product hairpin duplex allowing for RNA extension by 11 nucleotides. **d**, RNA elongation stalls at expected positions when the cognate NTP is withheld from the reaction. Extension to the end of the template is possible when MTP replaces either CTP or UTP in the presence of other substrate NTPs, showing that incorporation of M does not prevent RNA extension. Note that more efficient RNA extension is seen at higher NTP/MTP concentrations, also for MTP replacing UTP (not shown). Source data are provided as a Source Data file.

### RdRp uses NHC-containing templates to direct RNA mutagenesis

To investigate the templating properties of NHC, we prepared an M-containing RNA by solid-phase synthesis using the phosphoramidite building block M-PA, which we synthesized by the convertible nucleoside approach from a ribose-protected *O*^4^- chlorophenyluridine (**Fig. 3a**, Methods, **Extended Data Fig. 1, Supplementary information**). The presence of M in the obtained RNA and RNA purity were confirmed by denaturing HPLC and HR-ESI-MS (**Fig. 3b**). The M-containing RNA oligo was annealed with a fluorescently labeled product RNA such that the M nucleotide occupied the templating position +1 (**Fig. 3c, Supplementary Table 1**).

**Figure 3.**
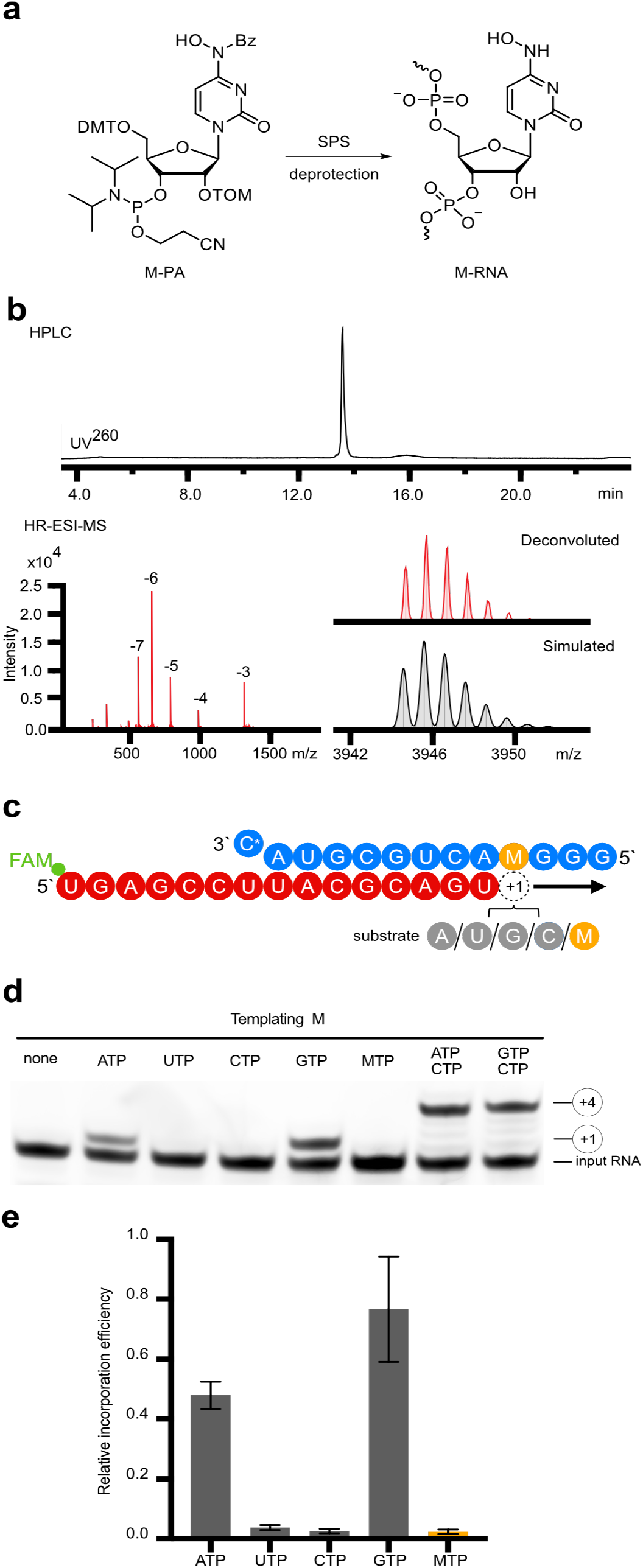
NHC can direct incorporation of G and A into RNA. **a**, Scheme of synthesis of RNA containing NHC monophosphate (M) at a defined position (M-RNA). **b**, Analysis of M-containing RNA by denaturing HPLC confirms homogeneity of the synthetic RNA (top). HR-ESI-MS proves the presence of NHC and absence of unmodified RNA (bottom). **c**, RNA template-product scaffold with M in the template position +1, where it is used by the RdRp to direct binding of the incoming NTP substrate. The 5’-end of the RNA product contains a FAM fluorescent label. C* in the 5’ end of the template is dideoxy-C (ddC). **d**, When present at position +1 of the template strand, M can direct the incorporation of G or A into nascent RNA, but not C or U. Source data are provided as a Source Data file. **e**, Quantification of the experiment in panel d after triplicate measurements. Incorporation efficiencies are calculated relative to C incorporation opposite of templating G.

Elongation assays showed that the M residue at the +1 position of the template strand directed incorporation of either G or A into nascent RNA, but not C or U (**Fig. 3d, e**). This can be explained by the formation of M–GTP or M–ATP base pairs in the RdRp active center. Consistent with this, thermal melting experiments with RNA duplexes containing M–G or M–A base pairs located at terminal or internal positions showed similar RNA duplex stabilities that were slightly lower than for duplexes containing a C–G base pair (**Extended Data Fig. 2, Supplementary Table 2**). Thus, when the RdRp uses RNA containing NHC monophosphate as a template, either the correct or the incorrect nucleotide is incorporated into the RNA product, and thus mutagenesis will occur.

### Structural basis of NHC-induced RNA mutagenesis

The above data indicate that the key aspect of the mutagenesis mechanism is the formation of stable M–G and M–A base pairs in the RdRp active center. To investigate this, we solved two structures of RdRp-RNA complexes that correspond to mutagenesis products after M-templated incorporation of either G or A (Methods). We formed RdRp-RNA complexes containing M in the template strand and either G or A at the 3’-end of the product strand. This was predicted to result in the formation of nascent M–G or M–A base pairs in position –1, which is occupied after successful M-templated nucleotide incorporation and RdRp translocation. We prepared RNA duplex scaffolds with M-containing oligonucleotides (**Extended Data Fig. 3**), formed RdRp-RNA scaffold complexes and subjected these to cryo-EM analysis as described^20^. We indeed obtained RdRp-RNA structures that contained either a M–A or a M–G base pair at position –1 (**Fig. 4, Table 1**). The structures showed an overall resolution of 3.3 Å and 3.2 Å, respectively, with the active center region resolved at ∼2.9 Å in both cases (**Extended Data Fig. 4**). As expected from the scaffold design, the structures showed the post-translocation state with a free NTP-binding site at position +1 (**Fig. 4b, c**). Comparison of the two structures with each other and with our original RdRp-RNA structure^20^ and remdesivir-containing RdRp-RNA structures^28^ did not reveal major differences, neither in the protein subunits nor in the nucleic acids, except that the protruding, second turn of RNA and the sliding poles of the nsp8 subunits were poorly ordered and not retained in the final model.

**Figure 4.**
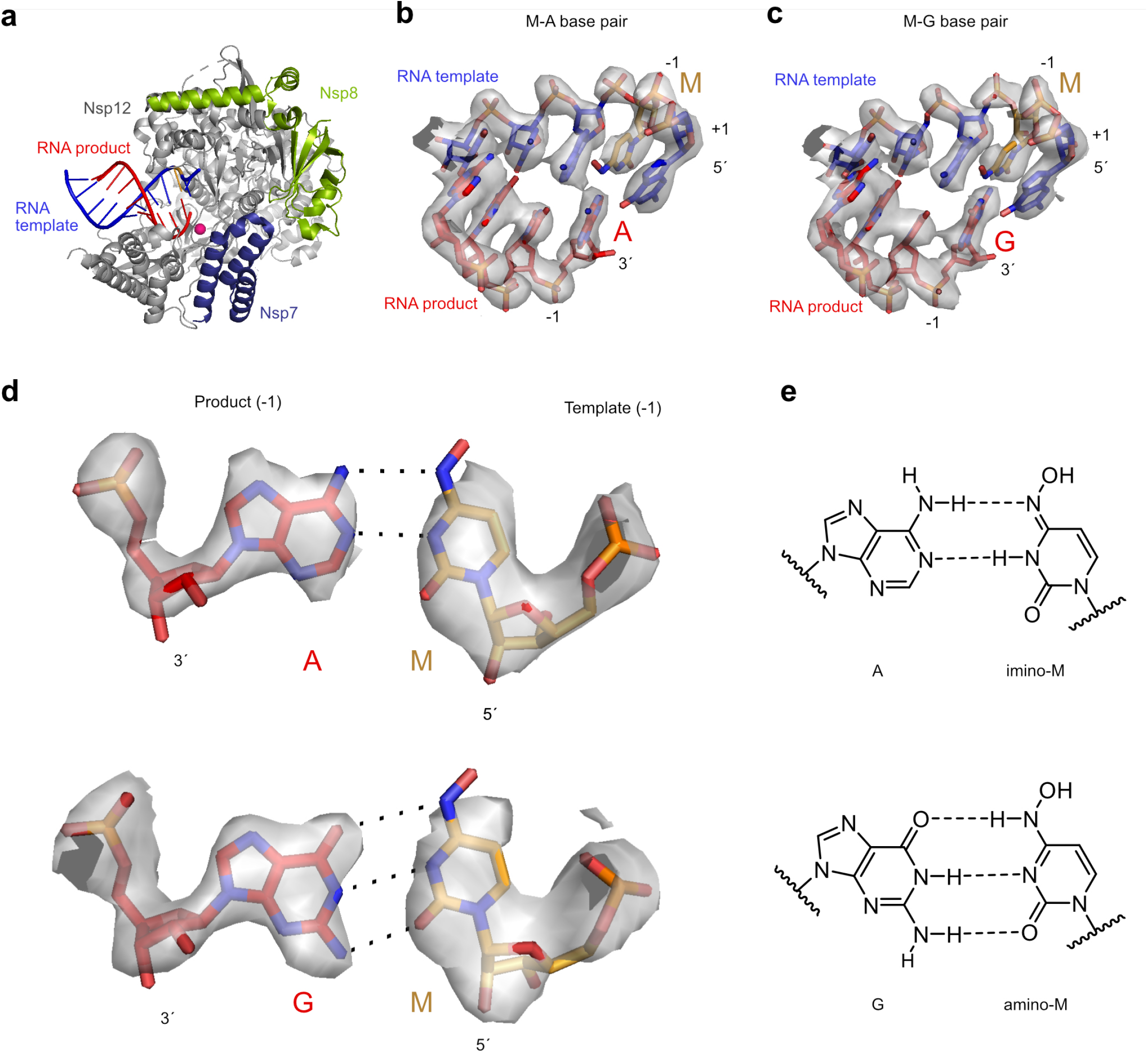
Structures of RdRp-RNA product complexes after NHC-induced mutagenesis. **a**, Overview of RdRp-RNA structure with M residue (orange) at position –1 in the RNA template strand. RdRp subunits nsp7, nsp8, and nsp12 are in dark blue, green and grey, respectively. The RNA template and product are in blue and red, respectively. The active site is indicated with a magenta sphere. Depicted is the structure containing the M–A base pair. **b**, RNA duplex containing the M–A base pair in the RdRp active center. The +1 position (templating nucleotide, NTP substrate site) and the –1 position (post-translocation position of nascent base pair) are indicated. **c**, RNA duplex containing the M–G base pair in the RdRp active center. **d**, Cryo-EM density for the nascent M–A (top) and M–G (bottom) base pairs in position –1, viewed along the RNA duplex axis in the direction of RNA translocation. **e**, M–A (top) and M–G (bottom) base pairing relies on different tautomeric forms of NHC45 as predicted^46^.

**Table 1:** Cryo-EM data collection, refinement and validation statistics.

|  | RdRp-RNA with M–A base pair<br>(EMD-XXXX)<br>(PDB XXXX) | RdRp-RNA with M–G base pair<br>(EMD-YYYYY)<br>(PDB YYYY) |
| --- | --- | --- |
| **Data collection and processing** |  |  |
| Magnification | 105,000 | 105,000 |
| Voltage (kV) | 300 | 300 |
| Electron exposure (e<sup>−</sup>/Å<sup>2</sup>) | 59.6 | 59.6 |
| Defocus range (μm) | 0.4-2.2 | 0.4-2.7 |
| Pixel size (Å) | 0.834 | 0.834 |
| Symmetry imposed | C1 | C1 |
| Initial particle images (no.) | 2,528,775 | 2,183,996 |
| Final particle images (no.) | 373,938 | 851,168 |
| Map resolution (Å) | 3.3 | 3.2 |
| FSC threshold | 0.143 | 0.143 |
| Map resolution range (Å) | 2.7-4.5 | 2.8-3.9 |
| **Refinement** |  |  |
| Initial model used (PDB code) | 7B3D | 7B3D |
| Model resolution (Å) | 2.9 | 2.9 |
| FSC threshold | 0.5 | 0.5 |
| Model resolution range (Å) | 2.7-4.5 | 2.8-3.9 |
| Map sharpening *B* factor (Å<sup>2</sup>) | -142 | -159 |
| Model composition |  |  |
| Non-hydrogen atoms | 8405 | 8406 |
| Protein residues | 991 | 991 |
| Ligands | 3 | 3 |
| *B* factors (Å<sup>2</sup>) |  |  |
| Protein | 66.68 | 61.32 |
| Ligand | 68.17 | 60.27 |
| R.m.s. deviations |  |  |
| Bond lengths (Å) | 0.004 | 0.005 |
| Bond angles (°) | 0.587 | 0.581 |
| Validation |  |  |
| MolProbity score | 1.52 | 1.56 |
| Clashscore | 7.14 | 8.23 |
| Poor rotamers (%) | 0.11 | 0.11 |
| Ramachandran plot |  |  |
| Favored (%) | 97.34 | 97.45 |
| Allowed (%) | 2.66 | 2.55 |
| Disallowed (%) | 0.00 | 0.00 |

The cryo-EM densities at the –1 position of the structures could readily be interpreted with the expected M–A or M–G base pairs (**Fig. 4b, c**). The densities were so detailed that we could clearly distinguish G and A bases (**Fig. 4d**), and were consistent with the proposed base pairing^46^ that is enabled by different tautomeric forms of NHC^45^ (**Fig. 4e**). However, the observed hydrogen bonding geometries were not optimal, possibly explaining our biochemical observations that suggest that M mimics C or U well, but not perfectly (**Figs. 1, 2**). These results represent the first direct visualization of NHC in a polymerase enzyme and show that stable M–G and M–A base pairs can be formed and accommodated in the RdRp active center, readily explaining our biochemical results.

## DISCUSSION

Our systematic biochemical analysis suggests a two-step model for the mechanism of molnupiravir-induced coronavirus RNA mutagenesis (**Fig. 5**). When the molnupiravir prodrug enters the cell, it is converted to NHC triphosphate (MTP), which can be used by the RdRp of SARS-CoV-2 as a substrate instead of CTP or UTP. Therefore, in a first step, the RdRp is predicted to frequently incorporate M instead of C or U when it uses the positive strand genomic RNA (+gRNA) as a template to synthesize negative strand genomic (–gRNA) and subgenomic RNA (–sgRNA). In a second step, the resulting M-containing RNA can be used as a template for the synthesis of +gRNA or positive strand subgenomic mRNA (+sgmRNA). The presence of M in the –gRNA then leads to mutations in the positive strand RNA products, which do not support formation of intact new viruses, as predicted by the ‘error catastrophe’ model^3,4,43^.

**Figure 5.**
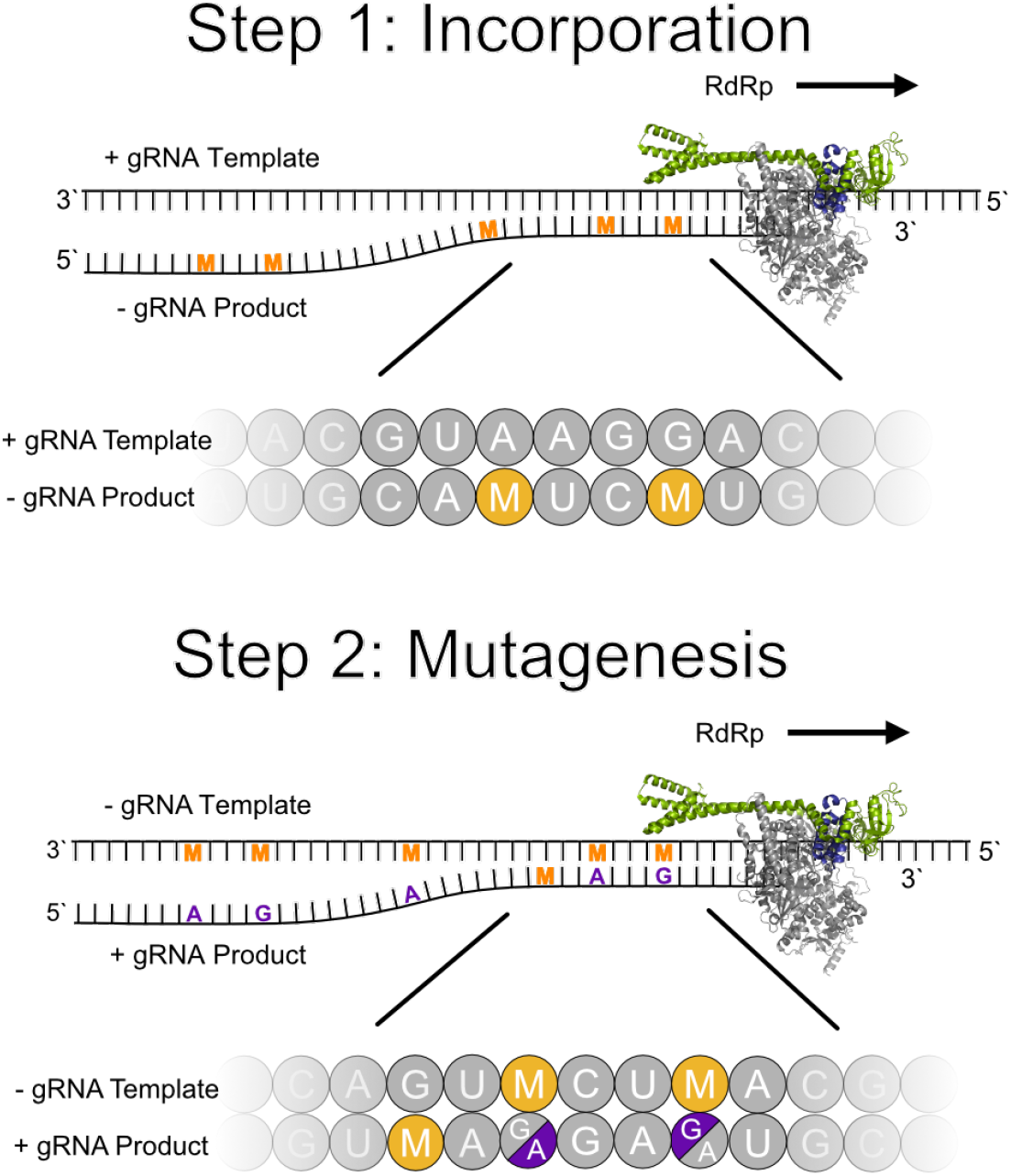
Two-step model of molnupiravir-induced RNA mutagenesis. In the presence of NTPs and MTP, M nucleotides can be incorporated by SARS-CoV-2 RdRp instead of C or U into the negative strand genomic (–gRNA) or subgenomic RNA (–sgRNA) during copying of the positive strand genomic RNA template (+gRNA). The obtained M-containing negative-strand RNAs can then be used as a template for the production of mutagenized +gRNA and positive strand subgenomic mRNA (+sgmRNA). These RNA products are predicted to be mutated and not to support formation of functional viruses. RNA of random sequence is shown with M and mutated residues indicated as orange and violet letters, respectively.

Our structural studies confirm the key aspect of this model, namely the formation of stable M–G and M–A base pairs in the RdRp active center. These base pairs mimic natural C–G and U–A base pairs, respectively, and do not impair RdRp progression. This antiviral mechanism is conceptually similar to the recently suggested mutagenesis mode of action of favipiravir^47,48^, but is entirely distinct from that of remdesivir, which impairs RdRp progression^28^. However, like remdesivir, molnupiravir escapes viral RNA proofreading because M incorporation and M-directed misincorporation are apparently not recognized by the viral exonuclease^23,24^. Such proofreading escape may also be due to the stability of the M– G and M–A base pairs that are predicted not to induce or favor backtracking of RdRp, which is likely required for exposing the RNA 3’-end to the proof-reading exonuclease^49,50^.

Finally, the two-step model can explain how molnupiravir or NHC monophosphate leads to RNA mutagenesis by polymerases of other viruses. For influenza, a possible two-step mutagenesis mechanism had been inferred from sequencing a molnupiravir-experienced virus population^3^. Also consistent with the model, MTP does not inhibit RNA synthesis by hepatitis C polymerase^40^ or the RdRp of respiratory syncytial virus^36^. Further, the reverse transcriptase of human immunodeficiency virus can incorporate G or A opposite of NHC that is located in the template^51^. The two-step mutagenesis model resides on the base-pairing properties of NHC that we structurally defined here and can explain why molnupiravir and NHC exhibit broad-spectrum antiviral activity against a wide variety of RNA viruses. In the future, it will be important to characterize the effects of molnupiravir and NHC on cellular polymerases and understand possible side effects of the antiviral agent.

## METHODS

No statistical methods were used to predetermine sample size. The experiments were not randomized, and the investigators were not blinded to allocation during experiments and outcome assessment.

### Protein preparation

Preparation of SARS-CoV-2 RdRp, composed of nsp12, nsp7 and two copies of the nsp8 subunits, was carried out as described^20^, with some modifications. Nsp12 protein was expressed from pFastBac vector 438C (Addgene #154759) in Hi5 insect cells. Production of the bacmid, V0 and V1 viruses was carried out as described^20^. 60 hours after transfection with corresponding V1 virus, the cells producing nsp12 with a 6xHis-MBP N-terminal tag were harvested by centrifugation (3000 rpm, 10 min at 4°C) and lysed by sonication in lysis buffer A1 (400 mM NaCl, 50 mM Na-HEPES pH 7.4, 10% (v/v) glycerol, 30 mM imidazole pH 8.0, 5 mM β-mercaptoethanol, 0.284 μg ml^−1^ leupeptin, 1.37 μg ml^−1^ pepstatin, 0.17 mg ml^−1^ PMSF and 0.33 mg ml^−1^benzamidine). Lysate was clarified by centrifugation at 74,766 g for 60 min and ultracentrifugation at 100,000 g at 4°C for 60 min, followed by filtration through 0.45 μm filter (Amicon Ultra Centrifugal Filter; Merck). The protein was bound to HisTrap HP prepacked columns (GE Healthcare) pre-equilibrated in lysis buffer, washed with high salt buffer B2 (1 M NaCl, 50 mM Na-HEPES pH 7.4, 10% (v/v) glycerol, 5 mM β-mercaptoethanol) followed by washing with buffer A1, and eluted with a gradient of 0-80% A1-B1 buffers over 30 column volumes (CV) (B1 buffer: 200 mM NaCl, 25 mM Na-HEPES pH 7.4, 10% (v/v) glycerol, 400 mM imidazole pH 8.0, 3 mM MgCl_2_ and 5 mM β-mercaptoethanol). Fractions containing nsp12 were pooled and the tag was cleaved with His-tagged TEV protease overnight during dialysis against buffer D (200 mM NaCl, 25 mM Na-HEPES pH 7.4, 10% (v/v) glycerol, 5 mM β-mercaptoethanol). The protein solution was applied to a HisTrap HP pre-packed column to remove uncleaved protein, tag and TEV protease. The flow-through containing nsp12 was further purified by ion exchange chromatography using Hi TRAP Q HP and SP HP pre-packed columns (GE Healthcare) equilibrated with buffer D. The unbound protein was concentrated and further purified via size exclusion chromatography on HiLoad S200 16/60 column (GE Healthcare) in buffer A2 (300 mM NaCl, 20 mM Na-HEPES pH 7.4, 10% (v/v) glycerol, 1 mM MgCl_2_, 1 mM TCEP). The peak contained monomeric fractions of nsp12 that were pooled, concentrated to 55 μM, aliquoted, flash-frozen in liquid nitrogen and stored at −80 °C.

Nsp7 and nsp8 were prepared as described^20^. Briefly, both proteins were expressed in *E. coli* BL21(DE3) RIL from the pET-derived vector 14-B (a gift from S. Gradia; Addgene 48308) in LB medium individually. Cells were grown to an optical density at 600 nm of 0.4 at 30 °C and protein expression was induced with 0.5 mM isopropyl β-D-1-thiogalactopyranoside at 18 °C for 16 h. After harvesting, cells were resuspended in lysis buffer A1 (400 mM NaCl, 50 mM Na-HEPES pH 7.4, 10% (v/v) glycerol, 30 mM imidazole pH 8.0, 5 mM β-mercaptoethanol, 0.284 μg ml^−1^ leupeptin, 1.37 μg ml^−1^pepstatin, 0.17 mg ml^−1^ PMSF and 0.33 mg ml^−1^ benzamidine). Nsp8 and nsp7 were purified separately using the same purification procedure. The cells were lysed using a French press and three cycles. Lysates were subsequently cleared by centrifugation (74,766 *g*, 4 °C, 30 min). The supernatant was applied to a HisTrap HP column (GE Healthcare), preequilibrated in lysis buffer. The column was washed with high-salt buffer B2 (1 M NaCl, 50 mM Na-HEPES pH 7.4, 10% (v/v) glycerol, 5 mM β-mercaptoethanol), and with buffer A1 (400 mM NaCl, 50 mM Na-HEPES pH 7.4, 10% (v/v) glycerol, 30 mM imidazole pH 8.0 and 5 mM β-mercaptoethanol). The protein sample was then eluted using a gradient of 0-80% B1 over 30 column volumes (CV) (B1 buffer: 300 mM NaCl, 50 mM Na-HEPES pH 7.4, 10% (v/v) glycerol, 400 mM imidazole pH 8.0, 3 mM MgCl2 and 5 mM β-mercaptoethanol). The protein sample was dialysed in buffer D (200 mM NaCl, 50 mM Na-HEPES pH 7.4, 10% (v/v) glycerol and 5 mM β-mercaptoethanol) in the presence of His-tagged TEV protease at 4 °C. The dialysed sample was subsequently applied to a HisTrap HP column (GE Healthcare), preequilibrated in A1 buffer. The flow-through that contained the untagged protein of interest was applied to an ion exchange chromatography HiTrap Q column (GE Healthcare). The unbound protein sample, containing nsp8 or nsp7, was concentrated using a MWCO 10,000 Amicon Ultra Centrifugal Filter (Merck) and applied to a HiLoad S200 16/600 (GE Healthcare) equilibrated in buffer A2 (300 mM NaCl, 20 mM Na-HEPES pH 7.4, 5% (v/v) glycerol, 1 mM TCEP). Peak fractions containing proteins were pooled, concentrated to 430 μM (nsp7) and 400 μM (nsp8), aliquoted, flash-frozen in liquid nitrogen and stored at −80 °C.

### RNA extension assays

All unmodified and 5’ FAM labelled RNA oligonucleotides (**Supplementary Table 1**) were purchased from Integrated DNA Technologies (IDT). MTP was from MedChemExpress and NTPs from ThermoScientific. The assay was performed as described^20^, except for the following changes. The final concentrations of nsp12, nsp8, nsp7 and RNA were 3 μM, 9 μM, 9 μM and 3 μM, respectively, except that 1 μM of RNA was used in the assays in figure panels 1f,g and 2d. The concentration of NTPs was 37.5 μM except for assays shown in figure panels 1f, 1g and 2d, where 4 μM was used. RNA in annealing buffer (50 mM NaCl, 10 mM Na-HEPES pH 7.5) was annealed by heating it to 75°C for 1 min and gradually cooling to 4°C. Annealed RNA and pre-mixed RdRp were incubated in reaction buffer (100 mM NaCl, 20 mM Na-HEPES pH 7.5, 5% (v/v) glycerol, 10 mM MgCl_2_, 5 mM β-mercaptoethanol) for 10 min at 30°C and the reactions were started by addition of NTPs. After 20 min incubation at 30°C the reactions were stopped with 2x stop buffer (7 M urea, 50 mM EDTA, 1x TBE buffer). RNA products were resolved on 20% denaturing polyacrylamide-urea gels in 0.5x TBE running buffer and visualized with a Typhoon 95000 FLA Imager (GE Healthcare Life Sciences).

### Preparation and analysis of NHC-containing RNA oligonucleotides

5’-*O*-DMT-2’-*O*-TOM-*O*^4^-chlorophenyluridine was prepared as described^52^, and converted to the corresponding *N*^4^-hydroxy-*N*^4^-benzoylcytidine 3’-(2-cyanoethyl) diisopropyl phosphoramidite (M-PA) in three steps. Details of the synthetic procedures are provided below and NMR spectra of isolated compounds are given in the Supplementary Information. RNA oligonucleotides were then prepared by solid-phase synthesis on CPG support (0.6 µmol scale) using 2’-*O*-TOM-protected ribonucleoside phosphoramidites (70 mM in CH_3_CN) and ethylthiotetrazol (ETT, 250 mM in CH_3_CN) as activator, with 4 min coupling time, as previously described^28,52^. The oligonucleotides were deprotected with 25% NH_4_OH/EtOH 3/1 at 55 °C for 6 h, followed by 1 M TBAF in THF for 12 h, and carefully purified by denaturing polyacrylamide gel electrophoresis to remove a minor fraction, in which NHC was converted to C during deprotection.

The purity and identity of the RNA oligonucleotides was analyzed by anion-exchange HPLC (Dionex DNAPac PA200, 2×250 mm, at 60 °C. Solvent A: 25 mM Tris-HCl (pH 8.0), 6 M Urea. Solvent B: 25 mM Tris-HCl (pH 8.0), 6 M Urea, 0.5 M NaClO_4_. Gradient: linear, 0–48% solvent B, 4% solvent B per 1 CV), and HR-ESI-MS (Bruker micrOTOF-Q III, negative ion mode, direct injection).

Thermal melting experiments were performed in 10 mM sodium phosphate buffer pH 7.0, 100 mM NaCl, at a RNA duplex concentration of 20 µM. Absorbance versus temperature profiles were recorded at 260 nm on a Varian Cary 100 spectrometer equipped with a Peltier temperature controller, at a heating rate of 0.5°C/min, for two heating and two cooling ramps between 10 and 90°C. Melting curves were normalized to the absorbance at 95°C, fitted to a two-state transition model with linearly sloping lower and upper baselines, and the melting temperatures were determined at the inflection point of the curves.

### Synthesis and characterization of M-PA

All reactions were performed under inert nitrogen atmosphere with dry solvents (CH_2_Cl_2_, CH_3_CN). For workup and purification distilled solvents (technical quality) were used. Column chromatography was performed on silica gel (Kieselgel 60, Merck) with a particle size of 0.040-0.063. TLC was performed on Alugram® aluminium sheets (Machery-Nagel, UV visualization, 254 nm). NMR spectra were recorded using Bruker Avance III (400 MHz) spectrometers. Chemical shifts (*δ*) are given in ppm, relative to the residual solvent signals as internal standards (CDCl_3_: ^1^H = 7.26, ^13^C = 77.16). Data are reported as: s = singlet, d = doublet, t = triplet, q = quartet, m = multiplet, br = broad; Coupling constants (*J*) are given in Hz. High-resolution (HR) electrospray ionization (ESI) mass spectra (MS) were recorded on a Bruker micrOTOF-Q III spectrometer. The detected mass-to-charge ratio (*m*/*z*) is given, as well as the calculated monoisotopic mass (**Supplementary Information, Extended Data Set 1; Extended Data Fig. 1**).

To synthesize compound **1** (5’-*O*-(4,4’-Dimethoxytrityl)-*N*^4^-hydroxy-2’-*O*-(triisopropyl-silyloxymethylcytidine), 5’-*O*-DMT-2’-*O*-TOM-*O*^4^- chlorophenyluridine (400 mg, 474 µmol, 1.0 eq.) was dissolved in anhydrous CH_3_CN (4 mL) under nitrogen atmosphere. DMAP (174 mg,1.42 mmol, 3.0 eq.) and NEt_3_ (331 µL, 2.37 mmol, 5.0 eq.) were added, followed by hydroxyl amine hydrochloride (165 mg, 2.37 mmol, 5.0 eq). After stirring for 21 h at ambient temperature, the reaction mixture was diluted with CH_2_Cl_2_ and washed with saturated aq. NaHCO_3_ (2x). The organic phase was dried over Na_2_SO_4_ and the solvent was removed under reduced pressure. The crude product was purified by column chromatography (*n*-hexane:EtOAc + 1% NEt_3_ 1:2 to 2:1) to yield the product (compound **1**, 230 mg, 308 µmol, 64%) as a colorless foam.

To synthesize compound **2** (*N*^4^-Benzoyl-5’-*O*-(4,4’-dimethoxytrityl) -*N* ^4^ - hydroxy - 2 ‘ - *O* - (t r i - isopropylsilyloxy)methylcytidine), a solution of compound **1** (200 mg, 267 µmol, 1.0 eq.) in anhydrous CH_2_Cl_2_ (4 mL) was treated with DMAP (65.3 mg, 535 µmol, 2.0 eq.) and NEt_3_ (149 µL, 1.07 mmol, 4.0 eq.) under nitrogen atmosphere. Benzoic anhydride (59.9 mg, 265 µmol, 0.99 eq.) was added in three portions within 3 h and the resulting reaction mixture was stirred for one more hour at ambient temperature. Volatiles were removed under reduced pressure. The crude residue was purified by column chromatography (*n*-hexane:EtOAc + 1% NEt_3_ 1:2) to yield compound **2** (183 mg, 215 µmol, 80%) as a colorless foam.

To synthesize compound **3**, (*N*^4^-Benzoyl-5’-*O*-(4,4’- dimethoxytrityl) - *N*^4^ - hydroxy - 2 ‘ - *O* - (triisopropylsilyloxy)methyl cytidine 3’-cyanoethyl-*N,N*-diisopropyl phosphoramidite), compound **2** (150 mg, 176 µmol, 1.0 eq.) was dissolved in anhydrous CH_2_Cl_2_ (2 mL) and cooled to 4°C. 2-Cyanoethyl *N,N,N′,N′*-tetraisopropyl phosphoramidite (67.3 µL, 212 µmol, 1.2 eq.) and 4,5-dicyanoimidazol (23 mg, 194 µmol, 1.1 eq.) were added in two portions within 1 hour. After one additional hour at 4°C, the reaction mixture was allowed to warm up to ambient temperature, and stirred for one more hour. Then, the solvent was evaporated and the crude residue was purified by column chromatography (*n*-hexane:EtOAc + 1% NEt_3_ 2:1 to 1:1) to yield compound **3** (75.0 mg, 71.3 µmol, 40%) as a colorless foam.

### Cryo-EM sample preparation and data collection

RNA scaffolds for structural studies were prepared by annealing of two RNA oligonucleotides because the length of the NHC-containing RNA was limited for technical reasons. The first RNA oligo was designed to form a template-product hybrid with a hairpin leaving a 10 nt product overhang at the 3’ end. A second oligo containing 8 nt complementary to the overhang was annealed to obtain an RNA template-product scaffold with a single nick at position -10 in the template RNA (**Extended Data Fig. 3, Table 1**). The second RNA oligo had NHC incorporated at position -1 and a short 5’
soverhang of three G nucleotides. RNA scaffolds for RdRp-RNA complex formation were prepared by mixing equimolar amounts of two RNA strands in annealing buffer (10 mM Na-HEPES pH 7.4, 50 mM NaCl) and heating to 75 °C, followed by step-wise cooling to 4°C. RNA sequences RdRp-RNA complexes were formed by mixing purified nsp12 (1.25 nmol) with an equimolar amount of annealed RNA scaffold and 3-fold molar excess of each nsp8 and nsp7. After 10 minutes of incubation at 30 °C, the mixture was applied to a Superdex 200 Increase 3.2/300 size exclusion chromatography column (GE Healthcare), equilibrated in complex buffer (20 mM Na-HEPES pH 7.4, 100 mM NaCl, 1 mM MgCl_2_, 1 mM TCEP) at 4 °C. Peak fractions corresponding to RdRp-RNA complex were pooled and diluted to 1.5 mg/ml. An additional 0.2 nmol of the annealed RNA scaffold were spiked into each sample prior to grid preparation. 3 µL of the concentrated RdRp-RNA complex were mixed with 0.5 µl of octyl ß-D-glucopyranoside (0.003% final concentration) and applied to freshly glow discharged R 2/1 holey carbon grids (Quantifoil). The grids were blotted for 7 seconds with blot force 5 using a Vitrobot MarkIV (Thermo Fischer Scientific) at 4 °C and 95 % humidity and plunge frozen in liquid ethane.

Cryo-EM data were collected with SerialEM^53^ on a Titan Krios transmission electron microscope (Thermo Fischer Scientific) operated at 300 keV. Inelastically scattered electrons were filtered out with a GIF Quantum energy filter (Gatan) using a slit width of 20 eV. Images were acquired using a K3 direct electron detector in counting mode (non-super resolution) at a nominal magnification of 105,000x, resulting in a calibrated pixel size of 0.834 Å/pixel. Images were exposed for a total of 2.0 seconds with a dose rate of 20.5 e^-^/px/s, resulting in a total dose of 59.6 e^-^/Å^2^ that was fractionated into 80 frames. Our previous cryo-EM analysis of SARS-CoV-2 RdRp-RNA complexes showed strong preferred particle orientation in ice^20^. To obtain more particle orientations, all data were collected with 30° stage tilt. Motion correction, CTF-estimation, and particle picking and extraction were performed using Warp^54^. A total of 11,060 and 10,230 movies were collected for M–A and M–G containing structures, respectively.

### Cryo-EM data processing and structural modeling

For the RdRp-RNA complex containing the M–A base pair, 2.5 million particles were extracted using Warp^54^ 1.0.9. Particles were imported to cryoSPARC^55^ 2.15 and subjected to 2D classification. 2D classes representing contamination or broken particles were selected and used for calculating two *ab initio* structures. All particles were then used for supervised 3D classification against five references, where four were originating from the *ab initio* reconstruction of contamination and broken particles and one was a previous RdRp-RNA complex structure (EMD-11995^28^). The class containing RNA-bound RdRp yielded ∼800,000 particles that were subjected to homogenous 3D refinement. The refined particles were then exported to RELION 3.1^56^ and focus-refined in 3D with an initial local angular sampling of 3.7° and a mask around RdRp that omitted the nsp8 sliding poles and the upstream, second RNA turn. To improve the quality of the density in the active site, particles were 3D classified without image alignment (T=4, four classes) using the same mask. The best class was focus-refined to an overall resolution of 3.3 Å. Local resolution was estimated with Relion 3.1 using a kernel size of 10 Å. For the RdRp-RNA complex containing the M–G base pair, 2.2 million particles were extracted using Warp^54^ 1.0.9. Further processing was as for the first complex, except that the class containing RNA-bound RdRp yielded 850,000 particles and the refinement resulted in a reconstruction at an overall resolution of 3.2 Å.

Atomic models were built using our previously published SARS-CoV-2 RdRp-RNA complex structure (PDB 7B3D^28^) as starting model. The model was first rigid-body fit into the density and then manually adjusted in Coot^57^. The protruding second RNA turn and the nsp8 extensions were removed from the model due to low quality of density for these regions. Restraints for molnupiravir monophosphate (M) were generated in phenix.elbow^58^ and the structures were refined using phenix.real_space_refine^59^ with secondary structure restraints. Model quality was assessed using MolProbity within Phenix^60^ which revealed excellent stereochemistry for both structural models (**Table 1**). Figures were prepared with PyMol (Schrödinger) and ChimeraX^61^.

## ACKNOWLEDGEMENTS

We thank U. Steuerwald for maintenance of the EM infrastructure. H.S.H. was supported by the Deutsche Forschungsgemeinschaft (SFB1190, FOR2848, EXC 2067/1-390729940). C.H. was supported by the DFG (SPP1784) and the ERC Consolidator Grant illumizymes (grant agreement No 682586). P.C. was supported by the Deutsche Forschungsgemeinschaft (SFB860, SPP2191, EXC 2067/1-390729940) and the ERC Advanced Investigator Grant CHROMATRANS (grant agreement No 693023).

## AUTHOR CONTRIBUTIONS

F.K. and J.S. designed and carried out biochemical experiments and analyzed biochemical data. C.S. synthesized and analyzed M-containing RNA oligonucleotides. F.K. and C.D. carried out cryo-EM data collection and processing. H.H. assisted with model building. C.H. designed and supervised RNA synthesis and analysis. P.C. designed and supervised research and wrote the manuscript, with input from all authors.

## DATA AVAILABILITY

The cryo-EM reconstructions and structure coordinates for the RdRp-RNA structures containing M–G or M–A base pairs were deposited with the Electron Microscopy Database (EMDB) under accession codes EMD-XXXXX and EMD-YYYYY and with the Protein Data Bank (PDB) under accession codes XXXX and YYYY, respectively. Source data are provided with this paper. Other data are available from corresponding authors upon reasonable request.

---

**Extended Data Fig. 1.**
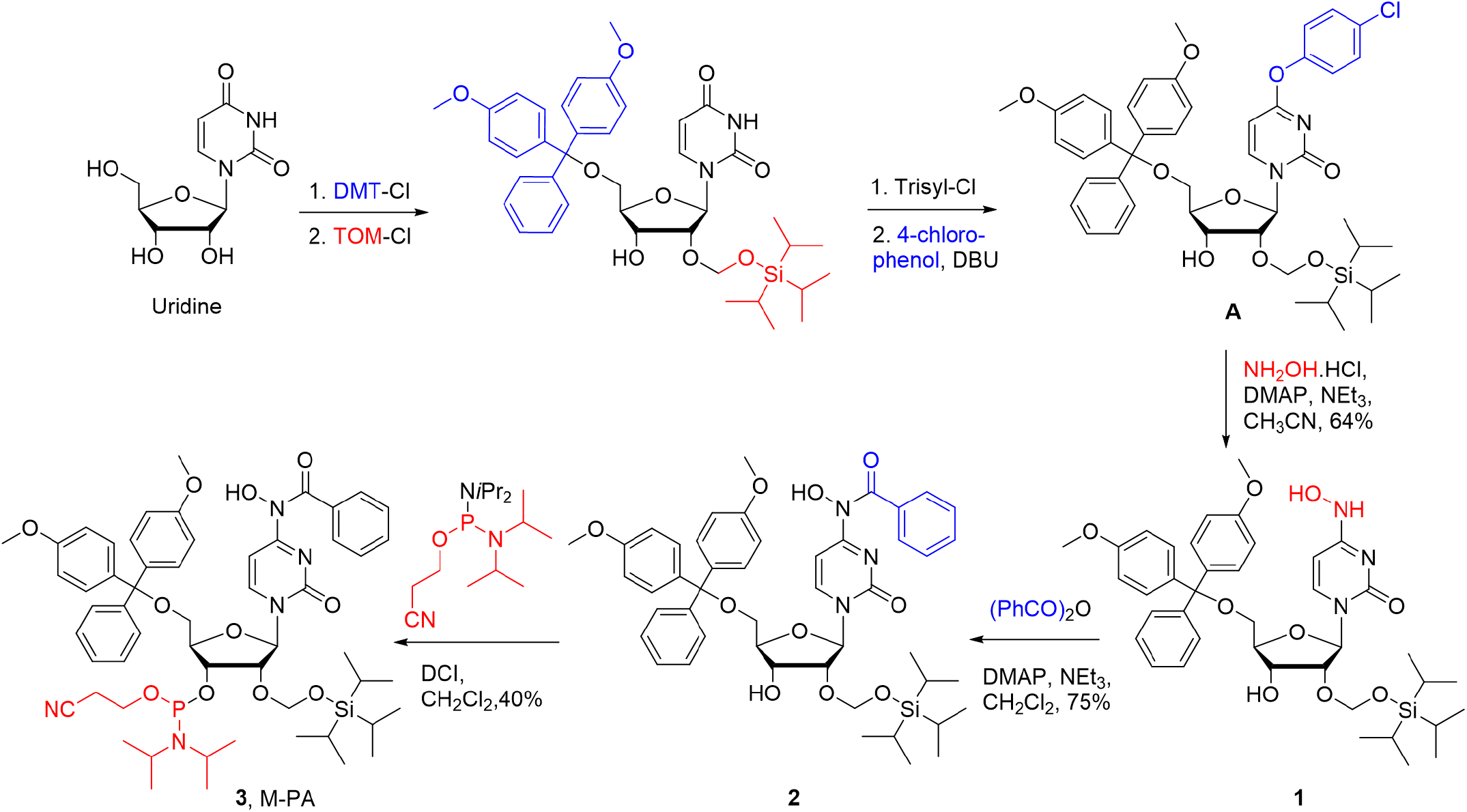
Synthesis of NHC-phosphoramidite 3 (M-PA). 5’-O-DMT-2’-O-TOM-O4-chlorophenyluridine (A) was synthesized from uridine as previously reported (L. Buttner, J. Seikowski, K. Wawrzyniak, A. Ochmann, C. Hobartner, Synthesis of spin-labeled riboswitch RNAs using convertible nucleosides and DNA-catalyzed RNA ligation. Bioorg Med Chem 21, 6171-6180 (2013)). The chlorophenol group was displaced by hydroxylamine to give new compound 1. After selective benzoyl protection at N4 with benzoic anhydride, compound 2 was converted to the phosphoramidite M-PA (3) using 2-cyanoethyl-N,N,N’,N’-tetraisopropylphosphorodiamidite and 4,5-dicyano-imidazol (DCI) in analogy to a previous report (J. Lu, L. Nan-Sheng, J. A. Piccirilli, Efficient Synthesis of N 4-Methyl- and N 4-Hydroxycytidine Phosphoramidites. Synthesis 16, 2708-2712 (2010)). DMT-Cl = 4,4’-dimethoxy-trityl chloride, TOM-Cl = triisopropylsilyloxymethyl chloride, DBU = 1,8-diazabicyclo [5.4.0]undec-7-ene, DMAP = 4-(N,N-dimethylamino)-pyridine.

**Extended Data Fig. 2.**
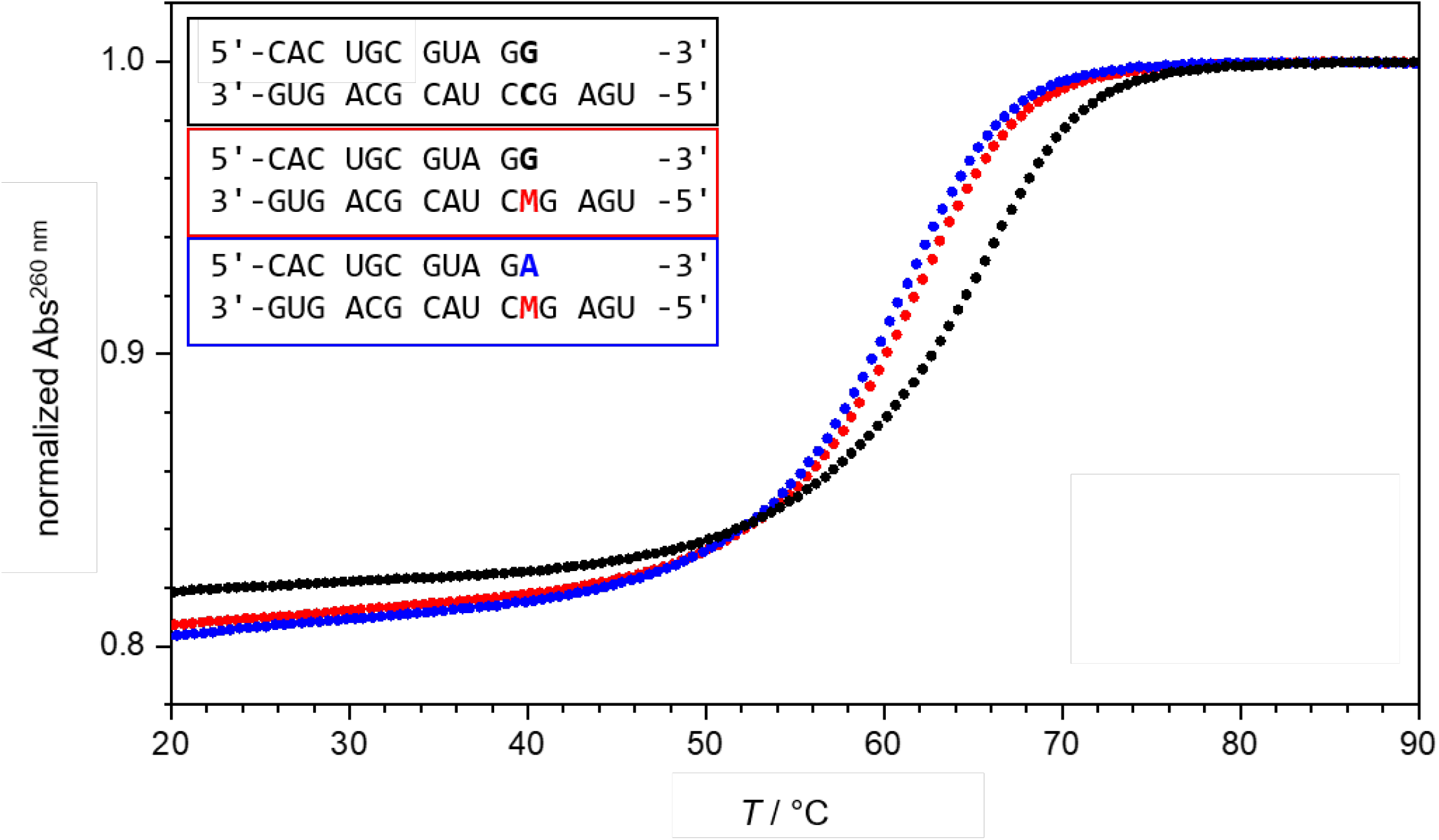
Melting curves for RNA duplexes containing M–G or M–A base pairs. UV thermal melting monitored at 260 nm for 20 µM duplexes (11 bp, 4 nt single-stranded overhang) in 100 mM NaCl, 10 mM Na-phosphate buffer pH 7.0. black: unmodified duplex C–G (Tm = 64.7°C), red terminal M–G (Tm = 61.2°C): blue: terminal M–A base pair (Tm = 60.6°C). See also Supplementary Table 2.

**Extended Data Fig. 3.**
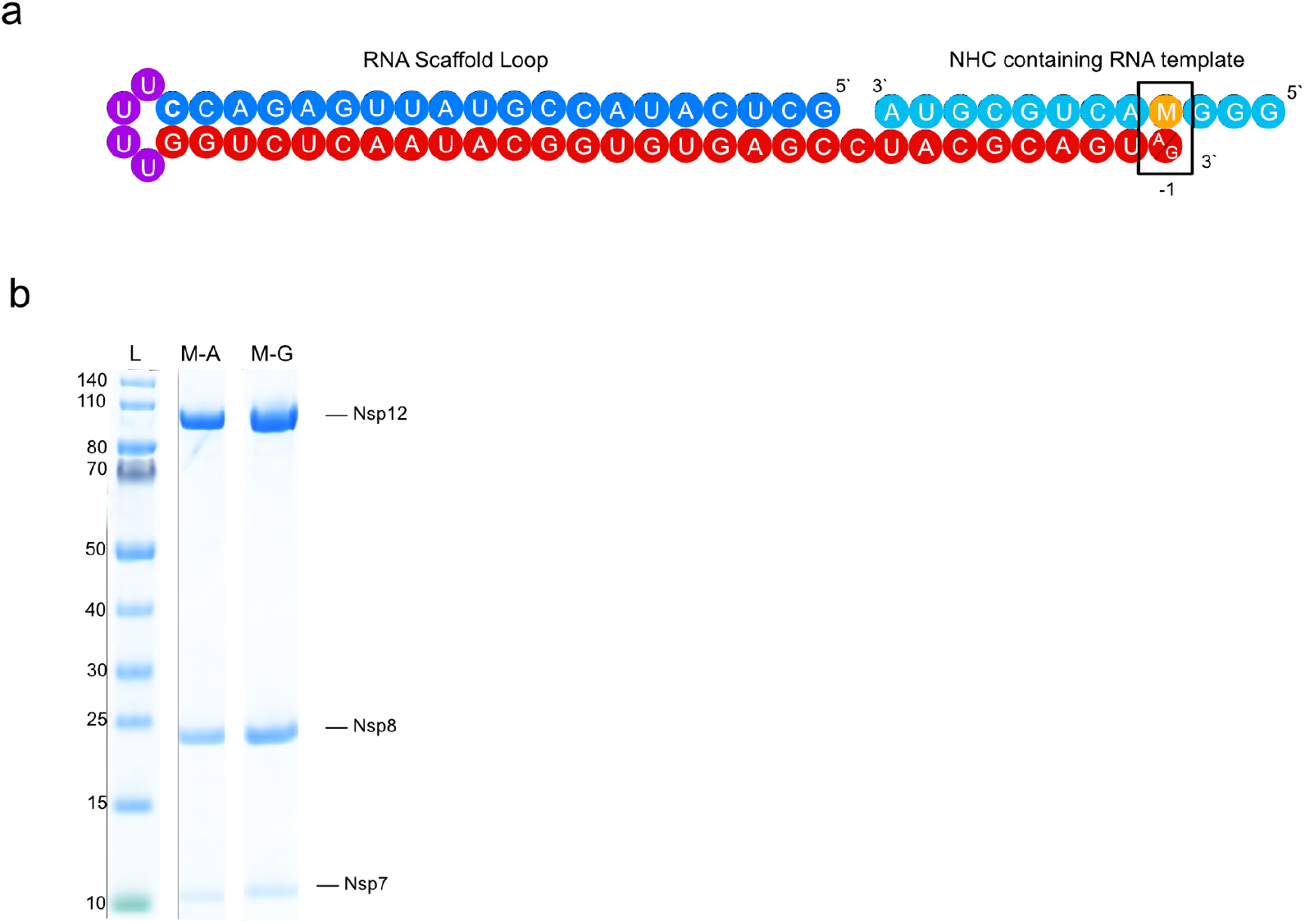
Protein preparation and RNA scaffold for structural studies. **a**, RNA scaffold was obtained by annealing a short M-containing oligonucleotide to a hairpin RNA duplex. **b**, SDS-PAGE of purified RdRp-RNA complexes used for cryo EM. Purified proteins were run on 4-12 % Bis-Tris SDS-PAGE gels in 1x MOPS buffer and stained with Coomassie Blue. M-A corresponds to the RdRp complex with RNA scaffold, where M in template base pairs to A. M-G corresponds to the RdRp complex with RNA scaffold, where M in template base pairs to G. L: PageRuler Prestained Protein Ladder (Thermo Scientific).

**Extended Data Fig. 4.**
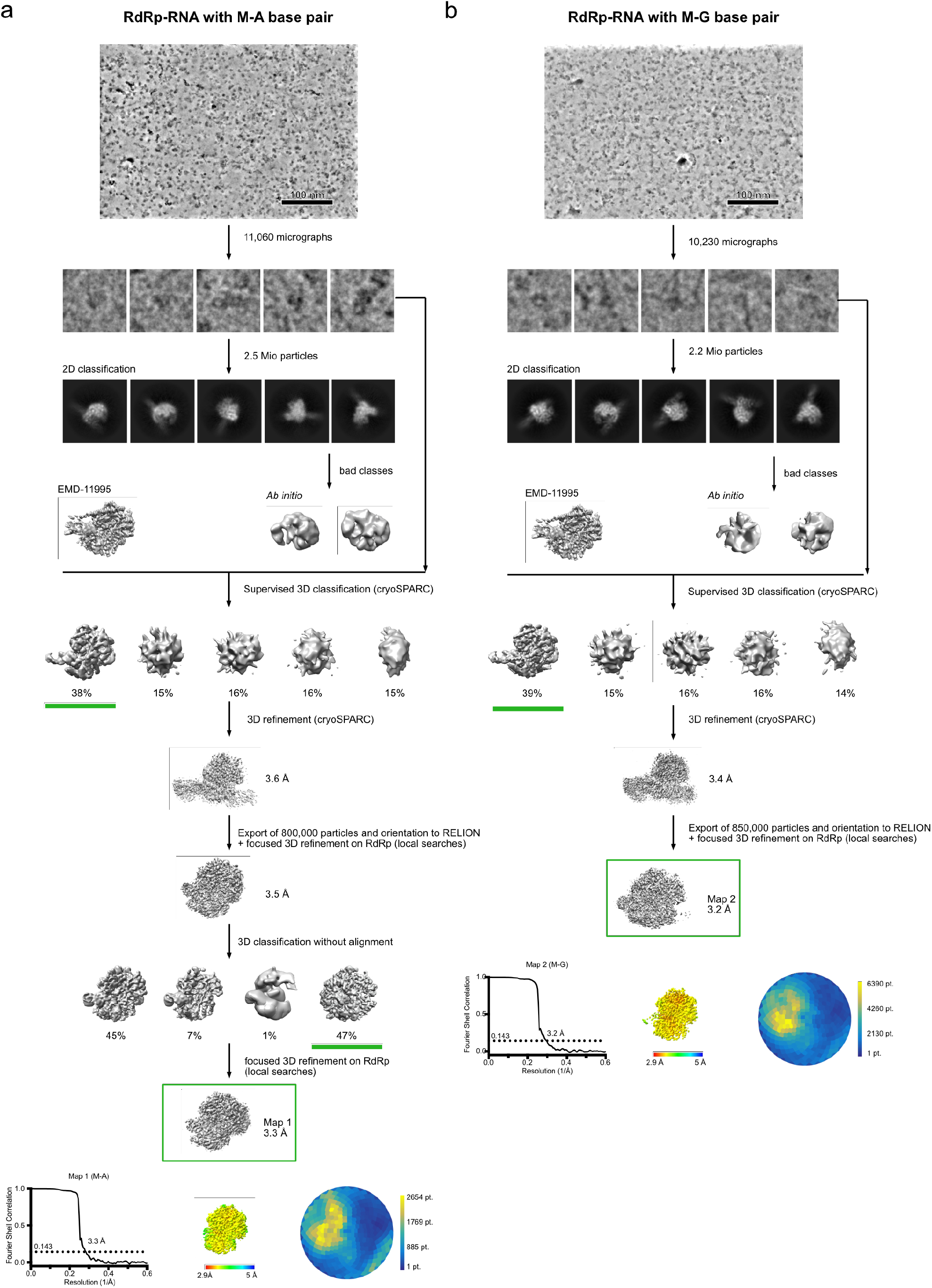
Cryo-EM sorting trees and quality of reconstructions. **a**, Cryo-EM sorting tree (top); local resolution, FSC plot and angular distribution of the final reconstruction (bottom) for M–A bp-containing RdRp-RNA structure. **b**, Cryo-EM sorting tree (top); local resolution, FSC plot and angular distribution of the final reconstruction (bottom) for M–G bp-containing RdRp-RNA structure.

